# Role of Sulphate Transporter *(PiSulT)* of Endophytic Fungus *Serendipita indica* in Plant Growth and Development

**DOI:** 10.1101/2020.01.07.897710

**Authors:** Om Prakash Narayan, Nidhi Verma, Abhimanyu Jogawat, Meenakshi Dua, Atul Kumar Johri

**Affiliations:** School of Life Sciences, Jawaharlal Nehru University, New Mehrauli Road, New Delhi-110067, India; School of Environmental Sciences, Jawaharlal Nehru University, New Mehrauli Road, New Delhi-110067, India

**Keywords:** *Serendipita indica*, sulphate transporter, maize, colonization, PiSulT

## Abstract

Sulfur is an important macronutrient required for the growth, development of plants and is a key component of many metabolic pathways. We have functionally characterized a high-affinity sulphate transporter (*PiSulT*) from an endophytic fungus *Serendipita indica*. The *PiSulT* belongs to the major facilitator superfamily (MFS) of membrane transporter. The *PiSulT* functionally complements the yeast sulphate transporter mutant HK14. *PiSulT* is a high-affinity sulphate transporter, having *Km* 15μM. We found enhanced expression of *PiSulT* in external fungal hyphae which helps the fungus in the acquisition of sulphate from the soil. When knockdown (KD)-*PiSulT*-*P.indica* colonized with the plant, it results in an 8-fold reduction in the transfer of sulphate to the colonized plants as compared to the plants colonized with the WT *S. indica*, which suggests that *PiSulT* is playing a role in sulphate transfer from soil to host plant. Further, plants colonized with the WT *S. indica* were found to be healthy in comparison to the plants colonized with the KD-*PiSulT*-*P.indica*. Additionally, *S. indica* colonization provides a positive effect on total sulfur content and on plant metabolites like sulfate ions and glutathione, particularly under low sulphate condition. We observed that the expression of sulfur assimilation pathway genes of *S. indica* and plant is dependent on the availability of sulphate and on the colonization with the plant. Our study highlights the importance of *PiSulT* in the improvement of sulfur nutrition of host plant particularly under low sulphate condition and in plant growth development. This study will open new vistas to use *S. indica* as a bio-fertilizer in the sulphate deficient field to improve crop production.

**One-Sentence Summary:** High-affinity sulphate transporter of *Serendipita indica* (*PiSulT*) transfer sulphate from soil to plant under low sulphate condition and improve plant growth and development.

## INTRODUCTION

Sulfur is an essential macronutrient for plant growth development and plays a fundamental role in metabolism. Sulfur is a structural component of protein disulfide bond formation, Fe-S group of electron transport chain, amino acids (cysteine and methionine), vitamins (biotin and thiamin), cofactors (S-adenosyl methionine) (Droux, 2004; Pilon-Smits and Pilon, 2007). Sulfur deficiency leads to a decrease in protein biosynthesis, chlorophyll content and eventually loss of crop yield (Sexton et al., 1997; Buchner et al., 2004; Lunde et al., 2008; Davidian and Kopriva, 2010). Sulfur contributes around 0.1% of the earth’s crust but it is very less accessible to living beings (Kertesz, 2000). Plants utilize sulfur primarily in its anionic form (SO4^2−^), which is generally available in very less amount in the soil. As sulphate is water-soluble, therefore it quickly loses from the soil by leaching (Eriksen and Askegaard, 2000; Buchner et al., 2004; Davidian and Kopriva, 2010). Under the condition of low sulfur availability in soil, a symbiotic association of an arbuscular mycorrhizal fungus (AMF) can help host plants to fetch sulfur from the soil. It has been established that a fungal partner helps host plant roots in nutrients uptake from nutrient-depleted soil rhizosphere, and in response, the fungal partner gets a carbon source from plants (Parniske, 2008). In this association, fungal nutrient transporter helps in nutrient transfer to host plant. It has been reported that the colonization of AMF *Glomus intraradices* can reduce sulfur starvation in plants like *Medicago truncatula* (Sieh et al., 2013). It has also been reported that AM symbiosis with the plant not only helps in nutrients uptake but also in the detoxification of metal contamination. For instance, *Rhizophagus irregularis* helps *M. truncatula* in the sulfur acquisition as well as in the chromium detoxification (Wu et al., 2018). It has been observed that sulphate transporter induced by both sulfur starvation and mycorrhiza formation show improved sulphate concentration in *Lotus japonicus* (Giovannetti et al., 2014). Arbuscular mycorrhizal colonization of *G. etunicatum, G. intraradices* with *Allium fistulosum* plants appears to make a substantial contribution to the sulfur status (Guo et al., 2007). It has been reported that AMF like *G. intraradices* helps in sulphate uptake and its translocation in the case of carrot especially under low sulphate condition (Allen and Shachar-Hill, 2009). Till date studies on sulphate transport in fungi have been limited to a few species such as *Saccharomyces cerevisiae, Neurospora crassa, Penicillium chrysogenum*, and *Aspergillus nidulans* (Breton and Surdin-Kerjan, 1977; Cherest et al., 1997; Marzluf, 1997; Van De Kamp et al., 1999; Van De Kamp et al., 2000; Piłsyk et al., 2007). However, due to the absence of a suitable transformation system in case of AMF, their sulphate transporter gene could not be genetically manipulated to improve sulfur uptake in plants colonized with AMF. Hence, mycorrhizal sulfur transfer to the host plant was poorly understood.

*S. indica* was isolated from the rhizosphere soils of the woody shrubs *Prosopis juliflora* and *Zizyphus nummularia* from the sandy desert soils of Rajasthan, northwest India (Verma et al., 1998). It has a typical pear-shaped chlamydospore and belongs to the newly formed order Sebacinales of Basidiomycota (Weiß et al., 2016). The size of *S. indica* genome is 24.97Mb having 1884 scaffolds and 2359 contigs (Zuccaro et al., 2011). *S. indica* has a wide range of host plants from bryophytes to angiosperms and monocots to dicots (Qiang et al., 2012). It colonizes the root of several economically important plants like rice, barley, wheat and showed mutualistic association with host plants (Jogawat et al., 2016). Unlike AMF, *S. indica* can be cultivated axenically and based on a well-established transformation system, studies have been conducted to understand the function of various genes in *S. indica* (Yadav, 2010; Akum et al., 2015). Association of *S. indica* with host plants provides several beneficial impacts to the host plant such as growth development, and it’s coping up with biotic and abiotic stresses (Waller et al., 2005; Kumar et al., 2009; Yadav, 2010; Johri et al., 2015; Jogawat et al., 2016; Narayan et al., 2017). Because of these qualities, *S. indica* has termed as plant probiotic (Aschheim et al., 2005). It has been reported that *S. indica* helps the colonized plants by the acquisition of nutrients such as phosphorus, magnesium, and iron from nutrient-deprived soil rhizosphere with the help of its nutrient transporters viz., phosphate, magnesium and iron transporter respectively (Yadav, 2010; Prasad et al., 2018, Verma et al 2019).

Characterization of sulphate transporter would provide insight into the regulation of sulphate uptake during symbiosis. In the present study, *PiSulT* has been, identified, isolated, functionally characterized and its role has been investigated in the sulphate transfer to the host plant. We demonstrate that *PiSulT* is essential for sulphate transfer to the host plant and helps in plant growth and development particularly under low sulphate condition. Additionally, this is the first report of a complete analysis of the regulation of sulphate assimilation pathway genes of *S. indica* and maize plants during colonization. Our results show that the biosynthetic steps are regulated at the levels of mRNA expression during adaptation under low sulphate condition. We suggest that the use of *S. indica* not only complements crop growth strategies but may also serve as a model system to study molecular mechanisms related to indirect uptake of sulphate by the plants and its regulation.

## RESULTS

### Identification and cloning of *PiSulT*

Our In-Silico analysis showed that putative sulphate transporter of S. indica belongs to PIRI_contig_0011 (Accession no CCA67103.1) in S. indica genome (Zuccaro et al., 2011). It has been annotated as probable sulphate permease, S. indica DSM11827. S. indica PiSulT shares 42% and 49% sequence identity and highest query coverage of 90% and 73% with S. cerevisiae sulphate transporters, Sul1, and Sul2 respectively. To amplify the putative PiSulT gene, total RNA isolated from the S. indica and further cDNA library was constructed. This cDNA was used as a template to amplify PiSulT with the help of gene-specific primers. The PCR amplified fragment was cloned into a pJET1.2 vector and further confirmed by sequencing and this fragment was sub-cloned into pYES2 yeast shuttle vector. We found PiSulT is 2292 bp long ORF. A deduced amino acid sequence of a putative PiSulT protein contains 763 amino acids and predicted polypeptide has a molecular weight of approx. 83.2 kDa. Sequences comparison showed that PiSulT has 7 exons and 6 introns (**Supplemental Figure 1, 2 & 3**).

### Phylogenetic and homology analysis

Our conserved domain analysis showed the presence of all important domains such as STAS (533-660), Sulphate_transp (164-455), Sulphate_tra_GLY (50-132), SUL1 (44-650) and PRK11660 (38-429) in PiSulT **(Supplemental Table 1).** It was also found that putative PiSulT has all the important motifs, domains, and sites that are essential for a protein to be defined as an MFS transporter. Importantly, we observed that relative spatial positions of STAS and catalytic domain in the case of *S. indica* PiSulT and *S. cerevisiae* Sul1 & Sul2 are the same. MultiAlin alignment of the deduced amino acid sequence of PiSulT with the amino acid sequence of sulphate transporter of a different kingdom and different group of fungus showed high and low consensus peptide sequence with highly conserved amino acids sequences at each position. The signature motif “**GLY**” of sulphate transporters is shown in the box and other conserved domains are shown in red shaded regions **(Supplemental Figure 4 and 5)**.

Phylogenetic analysis with diverse groups such as bacteria, insects, mammals, plants and fungi members were constructed to understand the position of putative *PiSulT* among fungi members and other groups. It was found that *PiSulT* is close to a member of Basidiomycota (Figure 1). *PiSulT* shares 42%, 33%, 30% and 29% sequence identity with sulphate transporter of *Saccharomyces cerevisiae*, *Homo sapiens, Drosophila melanogaster and Arabidopsis thaliana* respectively. Low sequence identity (27%) was observed with prokaryotic sulphate transporters such as *E. coli* and highest with fungal transporter (Table 1). Putative *PiSulT* showed the highest similarity to sulphate permease of fungus *Serendipita vermifera* (75%) and *Rhizoctonia solani* (65%) (**Supplemental Table 2**). It has more similarity with sulphate permease homolog protein of the member of Basidiomycota than the Ascomycota. (**Supplemental Table 3)**.

**Figure 1.**
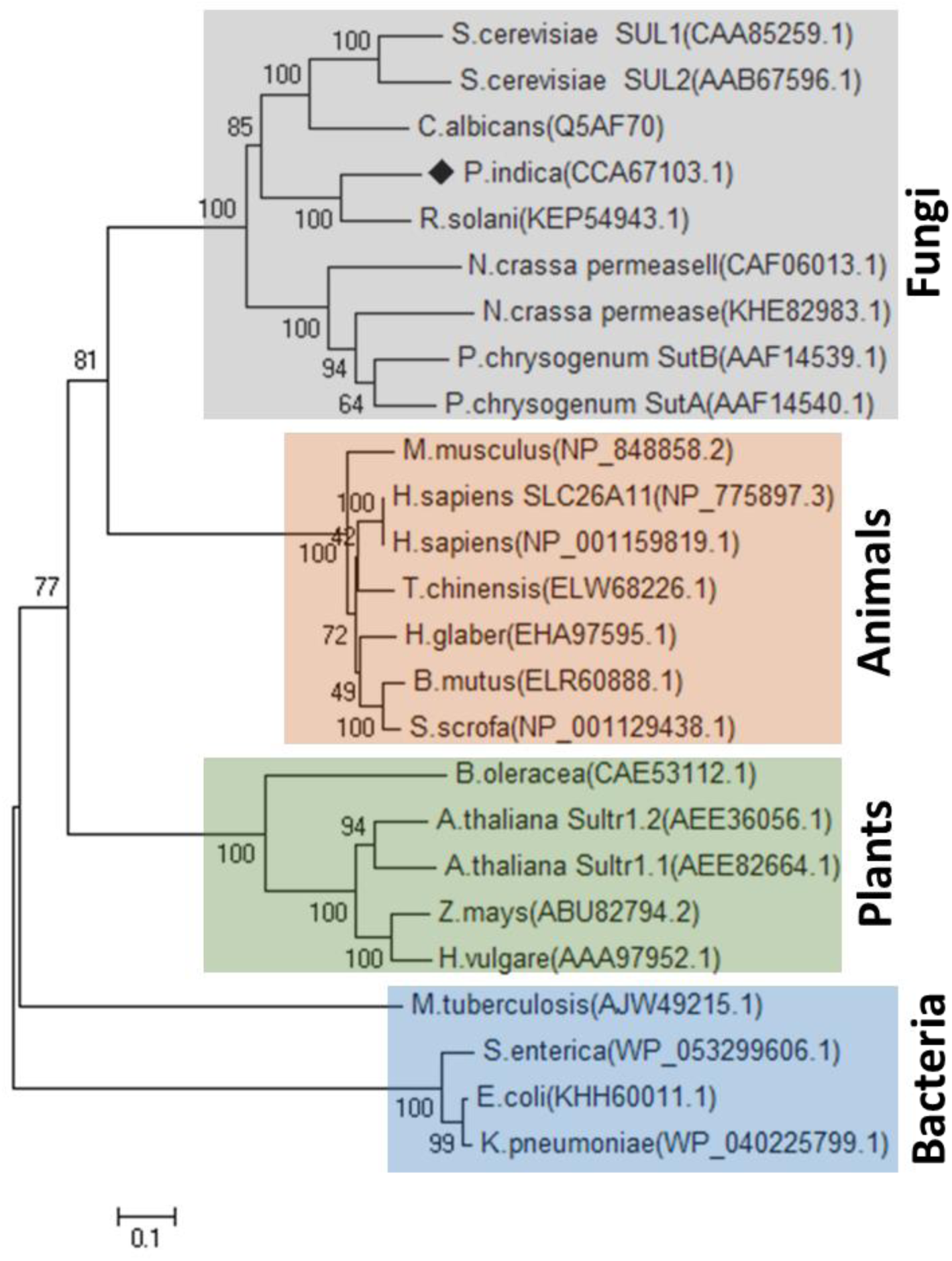
Phylogenetic tree of sulphate transporters. Phylogenetic tree of PiSulT with sulphate transporters of plants, animals, fungi, and prokaryotes. The tree was generated by MEGA-X software using Muscle for the alignment and the neighbor-joining method for the construction of the phylogeny. The bootstrap test was performed using 1000 replicates. The branch lengths are proportional to the phylogenetic distance. The distance scale showing genetic variation for the length of the scale (filled shape indicates *S. indica*). Value 0.1 showing distance scale, which represents the number of differences between sequences (e.g. 0.1 means 10 % differences between two sequences).

**Table 1.** Summary of amino acid identity (%) between *S. indica PiSulT* and other fungal, plant, animal, insect and prokaryotic sulphate transporters.

| **Organism Description** | **Sulphate Transporter**<br>**(no. of amino acids)** | **Gene Bank™**<br>**Accession No.** | **Identity with**<br>***PiSulT* %** |
| --- | --- | --- | --- |
| *S. indica* (Fungus) | PiSulT (763) | CCA67103.1 | 100 |
| *S. cerevisiae* (Fungus) | Sul1 (859) | AJP84964.1 | 42 |
| *H. sapiens* (Mammal) | SLC26A11 (606) | NP 001159819.1 | 33 |
| *D. melanogaster* (Insect) | Esp (623) | AAD53951.1 | 30 |
| *A. thaliana* (Plant) | SULTR1 (649) | OAO99615.1 | 29 |
| *E. coli* (Bacteria) | YchM (550) | SCQ14120.1 | 27 |

### *PiSulT* expresses more under low sulphate condition

To study the effect of sulphate concentration on *PiSulT* expression, *S. indica* culture were grown in MN medium containing different concentrations (1µM, 5µM, 10µM, 25µM, 50µM, 100µM, 1mM, and 10mM) of sulphate (Na_2_SO_4_). The *S. indica* was harvested at 1, 5, 10, and 15 days and RNA were isolated. The expression pattern of the *PiSulT* gene was determined by quantitative real-time PCR and semi-quantitative PCR. An increased expression level of *PiSulT* was observed at all the time points when sulphate was supplied at a concentration below100 µM (Figure 2A **&** 2B). This increased expression of *PiSulT* under low sulphate condition indicates the high-affinity nature of *PiSulT*.

**Figure 2.**
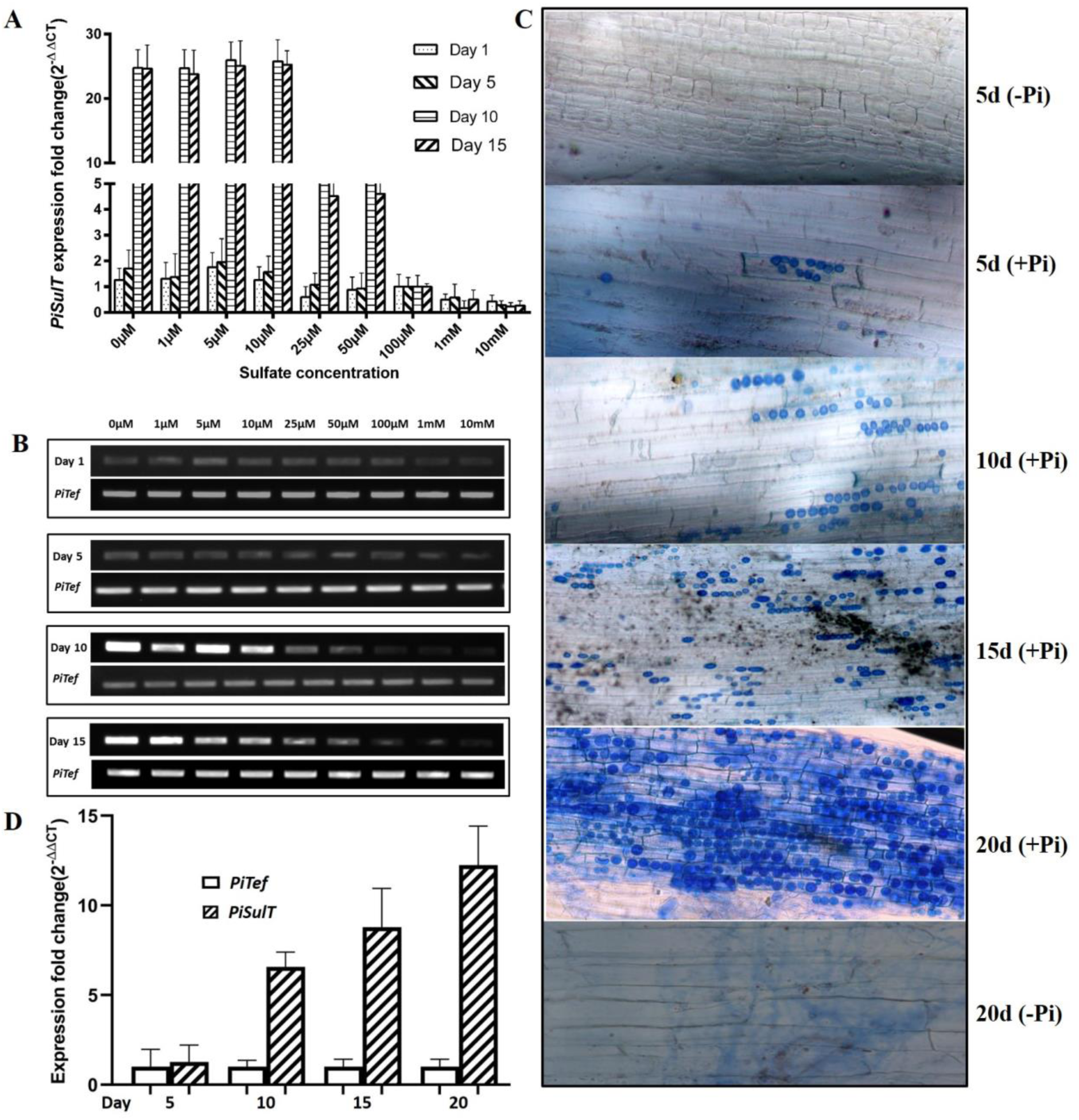
**Expression pattern of *PiSulT* in response to different concentrations of sulphate: A**. qRT-PCR of *PiSulT* gene isolated from *S. indica* grown in MN media containing the indicated different sulphate concentrations at 1, 5, 10, and 15 days. *PiTef* gene was used as a reference gene. **B**. Semi-quantitative RT-PCR of *PiSulT* gene isolated from *S. indica* under similar conditions. **Colonization of *S. indica* with maize root: C.** Trypan blue staining of maize plant roots showing the presence of intracellular chlamydospores of *S. indica* in the cortical cells at 5, 10, 15 and 20 days of colonization (+*PI*) (spores are shown in blue color) with the maize plant grown in sterile soil supplemented with the Hogland solution. 5 and 20-days plants without *S. indica* (-*PI*) were used as a negative control. **D**. **Expression of *PiSulT* during colonization with maize plant**: qRT-PCR showing the amplification of *PiSulT* transcripts from colonized maize plant roots at 5, 10, 15, and 20 days to confirm the presence of *S. indica* into plant root tissue. *PiTef* was used as a reference gene.

### Interaction of *S. indica* with maize plant

We found a maximum of 70 % colonization of the *S. indica* in the roots of the maize plant at the end of 20 days. Colonization was confirmed by histochemical analysis (Figure 2C). We found that as the colonization of *S. indica* increases in the root, a gradual increase in the expression of *PiSulT* also takes place (Figure 2C **and** 2D). Additionally, it was observed that colonization is associated with the developmental stage of the host tissue. *S. indica* showed strong colonization with newly formed lateral roots than tap roots. Heavy intercellular colonization in cortical tissue of differentiation and elongation zone was observed. No colonization was observed in the root tip meristem including root cap (**Supplemental Figure 6**). To validate this finding, we determined the amount of *S. indica* in different root zones by semi-quantitative PCR using *S. indica* genomic DNA as a template for the quantification of the *S. indica* translation elongation factor gene (*Tef*). A strong band intensity of *Tef* in the maturation zone was observed. However, a low-intensity band was observed in the apical zone (**Supplemental Figure 6i**).

### Complementation assay and growth analysis

Heterologous functional expression of *PiSulT* was analyzed in a yeast mutant cell of sulphate transporter HK14 (*Δsul1Δsul2*) by complementation and growth assay. For the complementation assay, *PiSulT* was cloned into a pYES2 yeast expression vector and transformed into HK14 mutant cells. For positive control, BY4742, a parental/WT strain of HK14 was used. This BY4742 cell contains both high-affinity sulphate transporter *sul1* and *sul2*. For negative control, mutant HK14 cells were transformed with empty vector pYES2. It is important to note that pYES2 has a galactose-inducible promoter, therefore, it can only express in the presence of galactose. Complemented mutant cells were tested to grow on the glucose and galactose supplemented media to confirm the controlled and regulated expression of a pYES2 vector having the *PiSulT* gene. It was observed that WT (BY4742) grew well on both glucose as well as galactose supplemented with the sulphate as they have a WT sulphate transporter gene. However, no growth was observed in the case of mutant transformed with the empty vector under both the conditions due to the lack of sulphate transporter. Mutant HK14 transformed with the *PiSulT* found to grow in the galactose only because of the induction of pYES2 vector by galactose and product of this gene restore the sulphate transport in HK14 cells (Figure 3Ai **and** 3Aii). Therefore, we conclude that *PiSulT* functionally complements the HK14 mutant. The growth pattern of all the above three types of cells at different concentrations of sulphate from low to high was also analyzed (Figure 3Bi **and** Bii). A similar growth pattern was observed in the case of WT and complemented strain, at all the concentrations of sulphate. However, in case of control (a mutant transformed with the empty vector), no growth was observed (Figure 3Biii). Nevertheless, the complemented strain showed the same growth pattern as of WT which indicates that the transformed *PiSulT* restore the sulphate transport activity in the mutant cells similar to that of WT.

**Figure 3.**
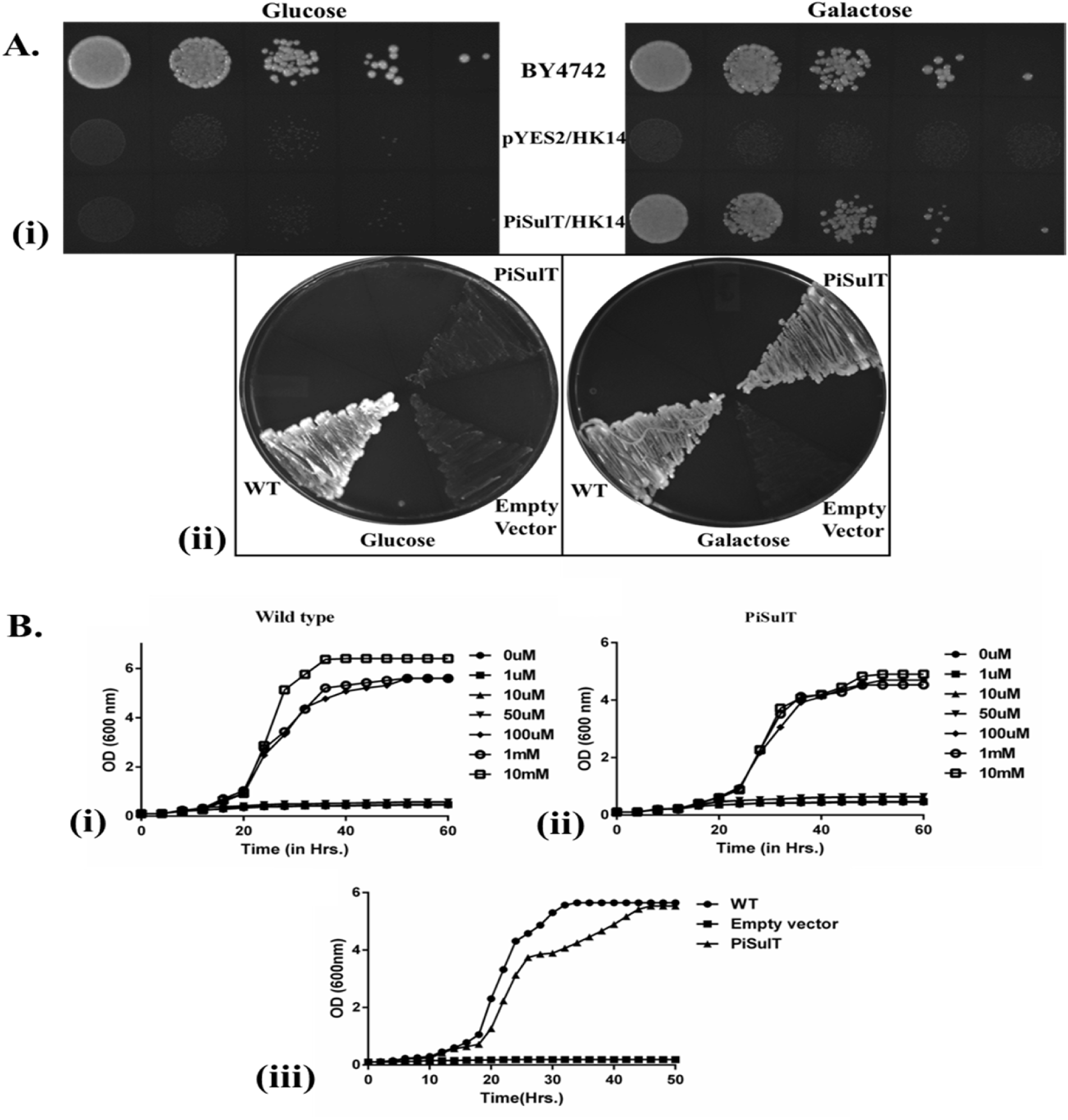
A. Drop test analysis for yeast sulphate transporter mutant HK 14 complementation. **(i)** Yeast cells were grown at 30°C on SD media containing 0.1 mM of sulphate as a sole source of sulfur in the presence of 2% glucose (non-inducing condition) and 2% galactose (inducing condition) as sole carbon source separately. Cells were suspended in sterile distilled water and cell density was adjusted to A_600_= 0.1, followed by serial dilutions of 1/10 (from left to right). Upper lane showing growth pattern of WT parent strain (BY4742) (used as a positive control), middle lane showing growth pattern of mutant transformed with empty vector (used as a negative control) and lower lane is showing growth pattern of mutant HK14 complemented with *PiSulT* **(ii)** Growth pattern (using streak method) of mutant HK14 complemented with *PiSulT*, WT and mutant transformed with the empty vector were shown under similar conditions mentioned above. **B. Growth pattern study**: **(i)** WT yeast (BY4742) **(ii)** mutant HK 14 transformed with *PiSulT*. In both, cases cells were starved for sulfur and then transferred to SD medium containing a different concentration of sulphate (as indicated) as the sole source of sulfur **(iii)** WT, mutant HK14 complemented with *PiSulT* and mutant HK 14 transformed with only vector (empty vector). In this case, cells were starved for sulfur and then transferred to medium containing 3mM concentration of sulphate as the sole source of sulfur.

### Chromate toxicity test to confirm the sulphate transport role of *PiSulT*

It has been established that chromate enters into the yeast cells through sulphate transporter. Transport of sulphate and chromate is a type of competitive transport, and it depends on the concentration of either of the substrates. To confirm the role of *PiSulT* in sulphate transport, we have performed the drop test. For this purpose, chromate toxicity was analyzed in the presence of different concentrations of sulphate. We observed that when the concentration of sulphate increases from 100µM to 1mM (with a constant concentration of chromate 20µM), there is a relief from chromate toxicity at higher concentrations of sulphate **(Supplemental Figure 7**). Further, WT, mutant and complemented HK14 were spotted on YNB plates containing an increasing concentration of chromate i.e. 40µm and 60µM. In the case of control, no chromate was used. Mutant strain, which does not have any sulphate transporter gene showed resistant to chromate and grew well. However, WT and complemented strain having sulphate transporter genes were susceptible to chromate toxicity and as a result, very little growth was observed (**Supplemental Figure 8**). This observation supports the sulphate transport nature of *PiSulT* gene.

### Sulphate uptake and kinetic analysis

For the functional characterization of *PiSulT*, the following sets were used **(a)** WT (BY4742) **(b)** mutant transformed with empty vector pYES2 (used as a control). **(c)** mutant transformed with *PiSulT* (Figure 4A). It was observed that 296 pmol of sulphate was transferred in the case of WT and 146 pmol in the case of complemented mutant. However, a negligible amount i.e., 0.2 pmol was found to be transferred by the mutant transformed with the empty vector (Figure 4A). The sulphate uptake by complemented HK14 cells expressing *PiSulT* follows typical Michaelis-Menten kinetics with an apparent *Km* of 15.0675±1.75 µM and *V*max value of 1.917±.063 pmol/min/A_650_ (Figure 4B). The *Km* of 8.2±1.38 µM and *V*max of 3.204±.041 pmol/min/A_650_ was observed in WT cells. To obtain the optimum pH value for the function of *PiSulT*, the mutant transformed with *PiSulT* was subjected to ^35^S-sulphate transport at different pH values ranging from pH 2 to 8. We found that sulphate transport activity of *PiSulT* is pH-dependent. The optimum value for sulphate transport by *PiSulT* was found to be pH 5 (Figure 4C).

**Figure 4.**
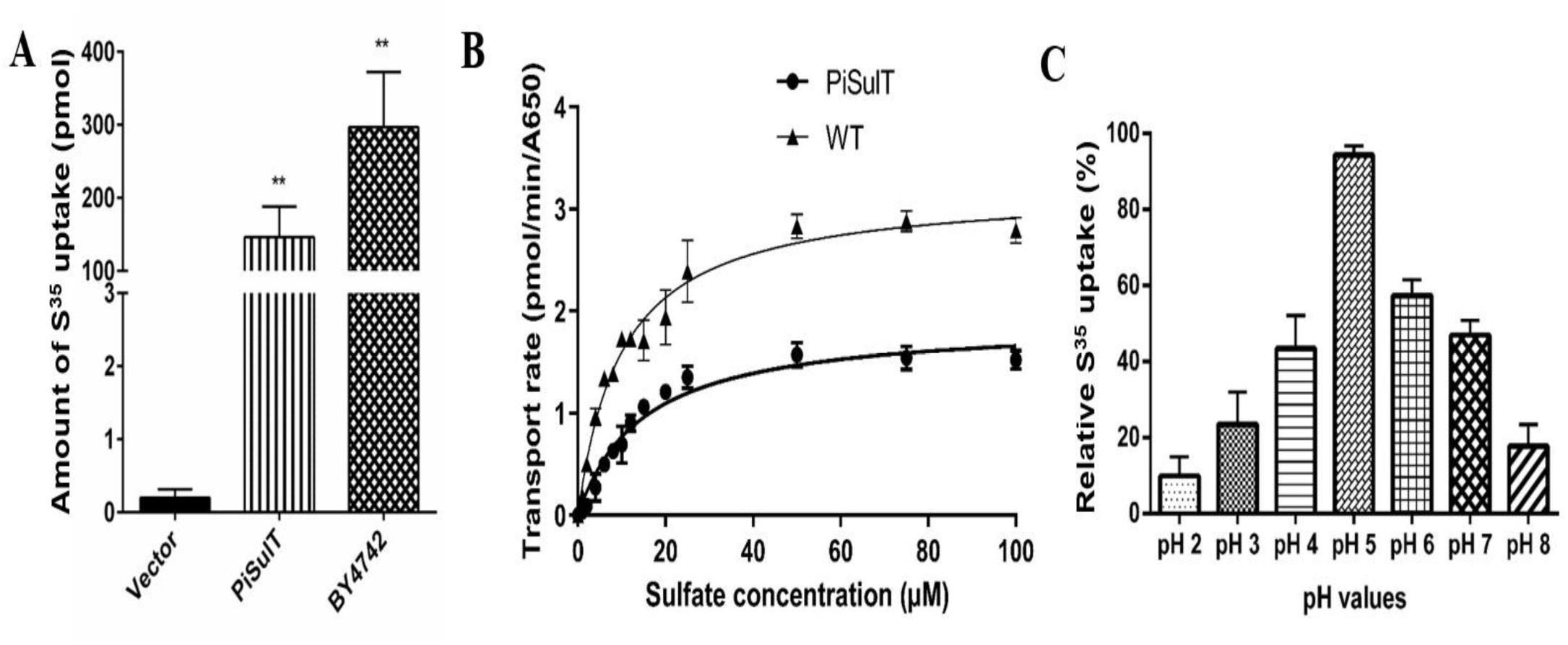
**Functional characterization of *PiSulT* gene using sulphate transporter mutant HK14 (*sul*1Δ&*sul*2Δ): A.** Sulphate uptake by yeast HK14 mutant cells transformed with empty vector (black), with *PiSulT* (vertical lines) and WT parent strain, BY4742, (crossed line) (used as a positive control). Means and standard errors of the means of the three replicate determinations consisting of three measurements **B**. Kinetics analysis of S^35^ uptake in a yeast mutant HK14 complemented with *PiSulT* and WT (BY4742). Nonlinear regression of sulphate uptake rate was measured after 4 minutes of time when cells transferred to media containing S^35^ labeled Na_2_SO_4_ in the presence of galactose at pH 5. The graph was calculated using Graph Pad Prism 6 software. The analysis was performed in triplicates and significance has been calculated using one-way ANOVA **C.** Determination of the optimum pH for sulphate uptake assay by *PiSulT*. The readings are relative to the negative control (HK 14 transformed with an empty vector).

### Role of *PiSulT* in sulphate transfer to host plant

To know the role of *PiSulT* in sulphate transfer to the host plant, knock-down sulphate transporter strain of *S. indica* (KD*-PiSulT-P.indica*) was developed by using RNA interference (RNAi). To knock-down *PiSulT* gene, we have used a special pRNAi vector having duel *S. indica* promoter PiTEF and PiGPD (**Supplemental Figure 9i**). The knockdown strain was selected on primary and secondary selection media containing Hygromycin as described in the method section (**Supplemental Figure 10A**). The expression of *PiSulT* in knock-down strain was analyzed by using qRT-PCR. It was found that *PiSulT* transcripts level was reduced in all obtained transformed colonies. However, in the case of TC1 transcript level was found to be very less **(Supplemental Figure 10B)**. The values obtained for *PiSulT* expression for WT (control), TC1, TC2, TC3 and TC4 were 1.0, 0.42, 0.68, 0.45 and 0.73-fold (∼60 % decrease in case of TC1) respectively relative to *PiTef*. Furthermore, TC1 showed the highest silencing of *PiSulT* expression as compared to other transformants and WT *S. indica*, hence selected for further experiments **(Supplemental Figure 10B)**. Further, the presence of the RNAi construct in knockdown *S. indica* was confirmed by PCR using hygromycin gene-specific primers. Amplification of a band was observed in all four transformants except WT *S. indica* (**Supplemental Figure 10C**). The siRNA accumulation was analyzed in the case of WT and KD-*PiSulT*-*S. indica.* We have observed the accumulation of siRNA in the case of KD-*PiSulT S. indica.* However, no detection of siRNA was observed in the case of WT (**Supplemental Figure 10D**). The growth of TC1 was also analyzed in KF broth and on KF agar plates. Both WT and TC1 colony grow in a similar fashion on KF media without Hygromycin. However, no growth of WT *S. indica* was observed as compared to TC1 when grown in KF supplemented with Hygromycin **(Supplemental Figure 11A and 11B)**.

The participation of *S. indica* in the transportation of sulphate from surroundings media to host plant was confirmed by using bi-compartment assay (**Supplemental Figure 12**). In the first set (set a) autoradiography revealed extensive labeling of maize plants by uptake of radiolabelled ^35^S in the case of WT *S. indica* (Figure 5Ai **and** 5Aii). The ^35^S was transferred to maize plants through the fungal mycelium and across the hyphal bridge between both compartments. Very little radioactivity was observed in the agar media of the second compartment confirming that the amount of ^35^S present in the maize plants was exclusively transferred by *S. indica* and not because of leaching by the fungus in the second compartment. In the case of set b, very less radioactivity was detected in maize plants colonized with KD-*PiSulT-P.indica* transformant, confirming direct role by *PiSulT* in sulphate transport to maize plants (Figure 5Bi **and** 5Bii). In the case of set c, no radioactivity was observed (Figure 5Ci **and** 5Cii) hence, the movement of ^35^S from one chamber to another was not due to diffusion but by the fungus only. We have observed that 362 pmol of sulphate was transported by WT *S. indica* to the host plant as compared to the 43 pmol in the case of KD-*PiSulT*-*S. indica* and this difference was found to be statistically significant (p<0.01) (Figure 5D). Colonization of both WT and KD-*PiSulT-S. indica* into maize plants was found to be similar in both the cases, i.e. 70 % at 20 dpi (Figure 5Aiii **and** 5Biii). It is important to note that in the colonized state, *S. indica* has external and internal hyphae. External hyphae ramify out of the colonized root and internal hyphae penetrate the root cortex. To determine the expression of *PiSulT* in internal and external hyphae, transcript abundance was measured by using relative quantitative RT-PCR. It was observed that *PiSulT* transcripts were 2.25 folds higher in the external hyphae as compared to internal hyphae and this was found to be statistically significant (P<0.05) (Figure 5E).

**Figure 5.**
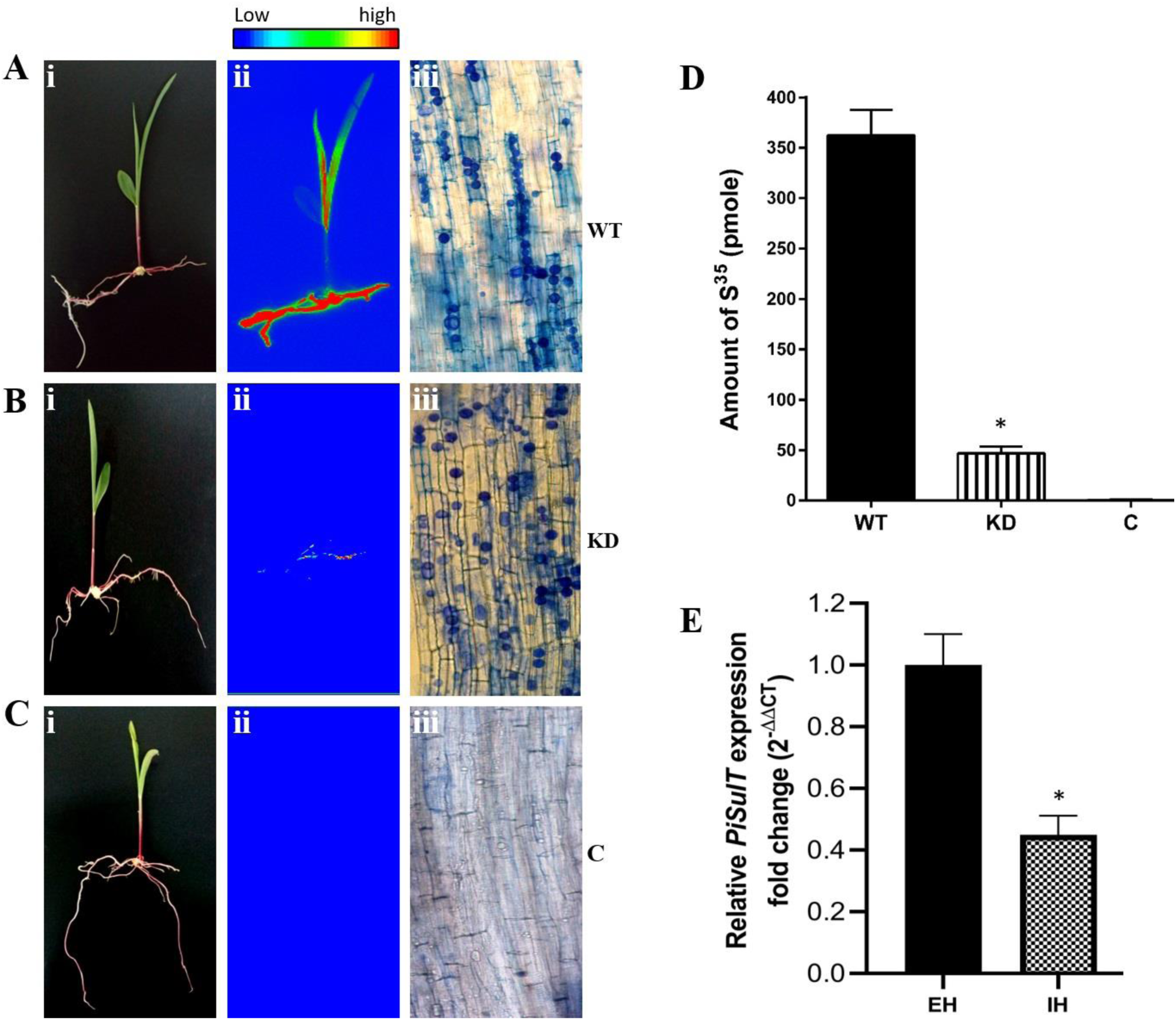
Transport of sulphate to maize plants by *S. indica* carried out in the bi-compartment Petri dish culture system. Radioactivity incorporated into plants was demonstrated by autoradiography. Radioactivity count intensities are shown in the false color code (vertical bar, low to high). **Panels. A.** Maize plants were colonized with WT *S. indica* designated as WT, **B.** Maize plants were colonized with KD-*PiSulT*-*S. indica* designated as KD; **C.** Maize plants were grown alone without *S. indica* designated as C. **(i),** whole maize plant before autoradiography; (**ii),** false-color autoradiograph of the maize plant obtained after 12 h of exposure of the maize plant; and (**iii)**, microscopic view of a sample of plant root showing colonization and non-colonization. **D.** The content of transferred sulphate: Amount of ^35^S transferred to the maize plant components by *S. indica*. Radioactivity was measured three times independently (the number of transformants used was *n*=3). The mean S.D. of three independent measurements is shown. * indicate a significant difference (p<0.01). **E. Spatial expression of *PiSulT* during colonized condition**. Expression of *PiSulT* in the external (EH) and internal hyphae (IH) of *S. indica* during colonization with maize plant. Relative fold change in IH was compared with EH. *Tef* gene was used as an internal control. The values obtained for *PiSulT* expression for EH and IH were 1 and 0.449-fold respectively. The means S.D. of three independent determinations is presented. * indicate significant difference from EH (p<0.01).

### Plant responses to sulfur deficiency and the role of *PiSulT* in sulphate nutritional improvements of the host plant

The role of *PiSulT* in sulphate nutritional improvement of the host plant was analyzed. Plants colonized with WT *S. indica* were found to be healthy as compared to the non-colonized plants and plants colonized with the KD*-PiSulT-S. indica*, grown under low sulphate condition (Figure 6A). Under similar conditions, biomass (fresh weight) of maize plants colonized with WT *S. indica* was found to be 1.8 fold and 2.4 fold higher and sulphate contents were found to be 1.6 and 2.3 fold in comparison of plants colonized with the KD-*PiSulT-P.indica* and non-colonized plants respectively and this was found to be statistically significant (p≤0.001) (Figure 6B **and** 6C). In a separate study to know the performance of the *S. indica* in the growth promotion activity of the plants, the plant responses under low sulphate condition (10µM) and sulphate-rich (10mM) conditions were chacked. For this purpose, four sets were prepared: **(1)** maize plants grown under low sulphate condition and treated with autoclaved macerated fungal mycelium (served as a control) **(2)** maize plants colonized with WT *S. indica* and grown at low sulphate condition, **(3)** maize plants grown under high sulphate condition and treated with macerated fungal mycelium (served as a control) **(4)** maize plants colonized with WT *S. indica* and grown at high sulphate condition. All four experimental sets were grown in acid-washed sand fertilized with a modified 0.5X Hoagland solution **(Hoagland and Arnon, 1950**) containing respective sulphate concentrations. After 4 weeks, plants were harvested, and fresh weight was measured. We observed that biomass (in terms of fresh weight) of maize plants colonized with WT *S. indica* was 1.2 fold higher when grown at sulphate-rich condition, whereas it was 2.3 fold higher in the case of maize plants colonized with WT *S. indica* and grown under low sulphate condition in comparison to their respective controls (p≤ 0.05) (Figure 6D). The total sulfur content was estimated in such conditions and, it was observed 2.3 and 1.6 fold high under similar conditions (Figure 6E). We also analyzed the content of plant metabolites such as glutathione and sulphate ions in plants under similar conditions during colonization. Glutathione content was found to be 1.8 and 0.8 fold and sulphate ions were found to be 1.5 and 0.6 fold higher under similar conditions (Figure 7 A & B).

**Figure 6.**
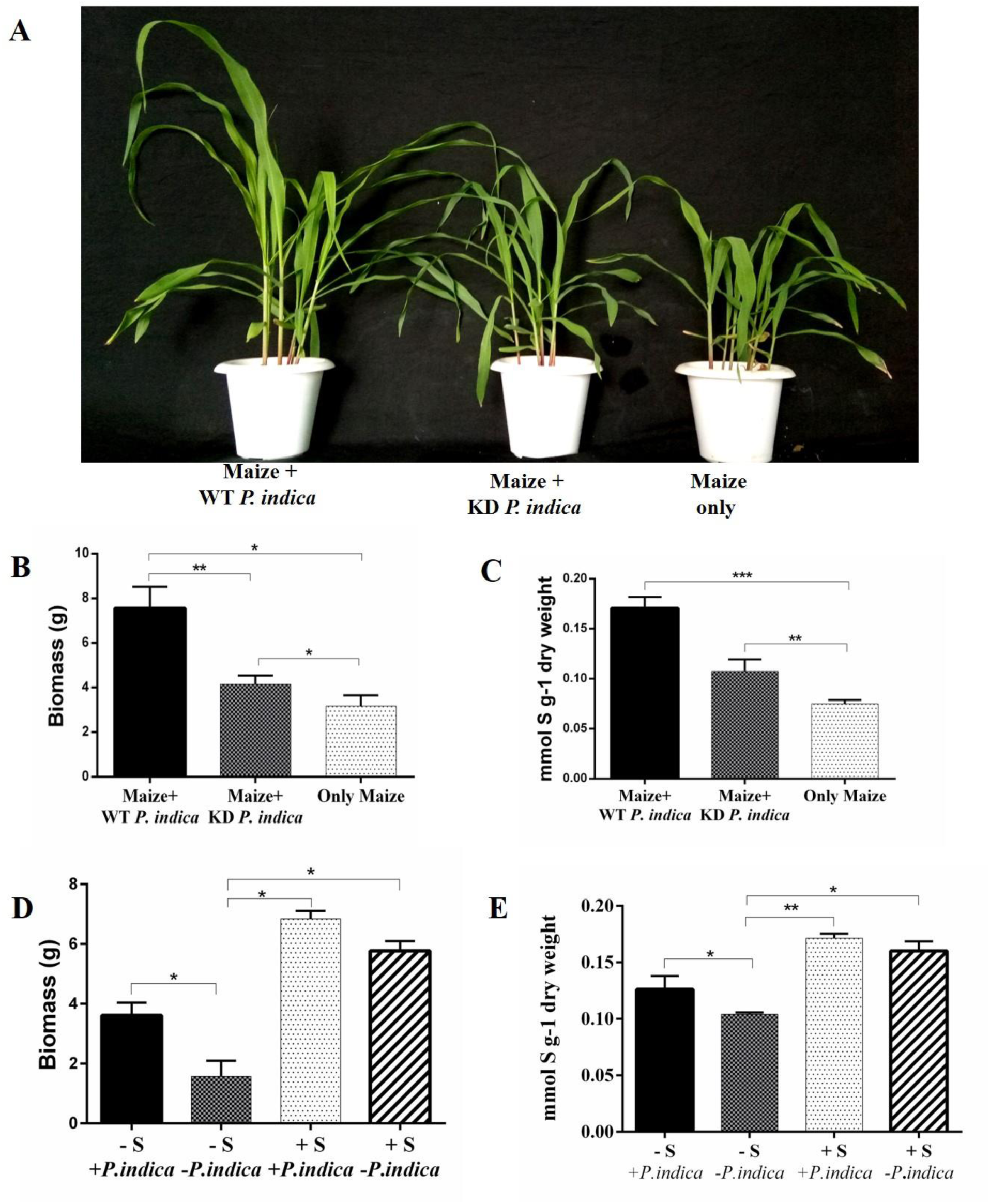
The impact of *PiSulT* on plant health and development and on sulphate nutritional enrichment. **A.** Maize plant colonized with WT *S. indica* showing improved growth as compared to maize plants colonized with KD-*PiSulT*-*S. indica* and maize plants without *S. indica.* **B.** Biomass study of maize plants colonized with WT *S. indica* showing improved growth as compared to maize plants colonized with KD-*PiSulT*-*S. indica* and maize plants without *S. indica* (control). **C**. Sulphate content. **D.** The impact of *S. indica* on host plant under high or low sulphate concentration. Total biomass (fresh weight) of maize plants grown under low (10uM) and high (10mM) sulphate conditions with or without *S. indica* colonization. **E.** Sulphate content. (-S= low sulphate, +S= high sulphate +Pi= plant colonized with *S. indica*, –Pi= non-colonized plant). Different asterisk indicates significant differences, * significant (p≤0.05), **(p≤0.01), ***highly significant (p≤0.001). Tukey test was used to check the significance.

**Figure 7.**
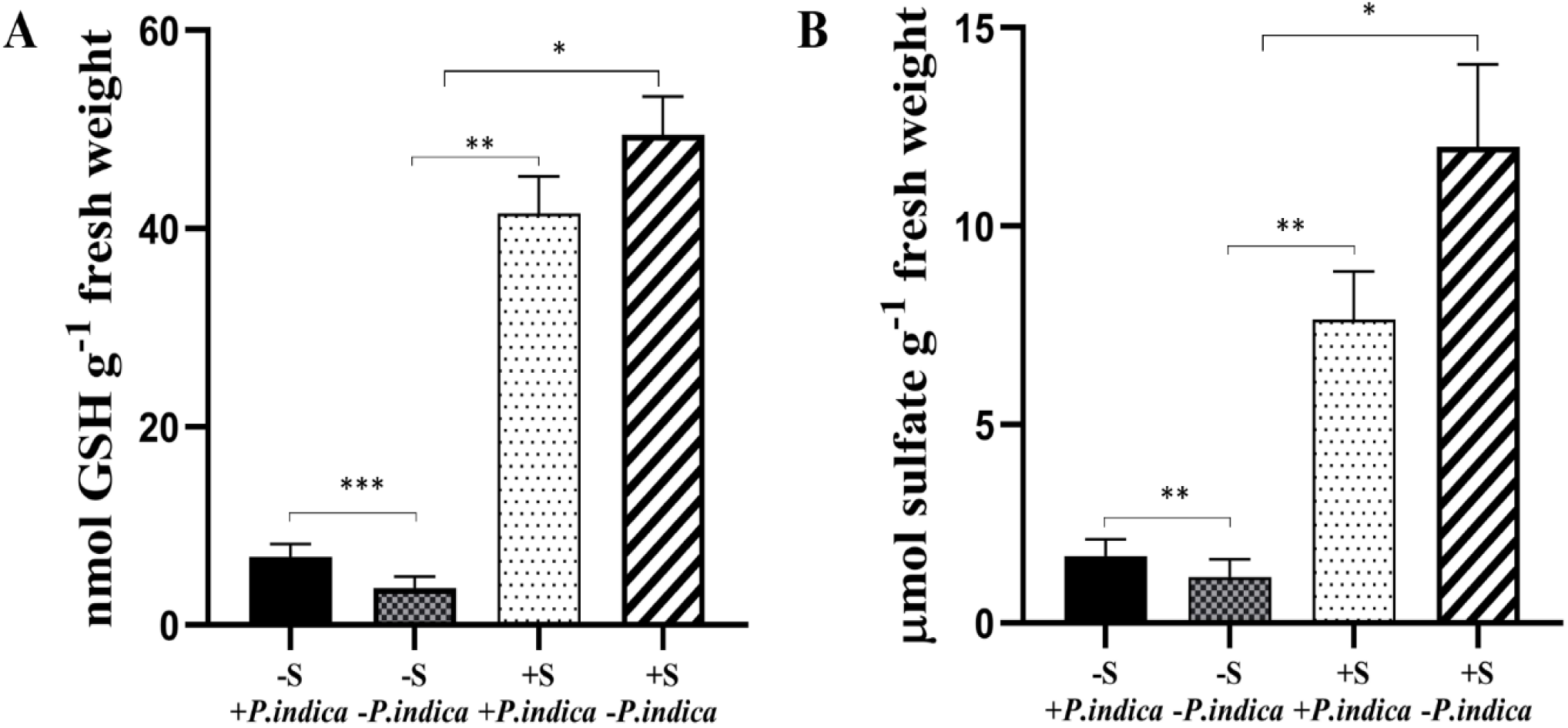
The impact of *PiSulT* on the improvement of plant metabolite and sulphate nutrition. Measurement of glutathione (GSH) content (**A)** and sulphate content (**B**) in maize plants grown under low (10uM) and high (10mM) sulphate conditions with or without *S. indica* colonization (-S= low sulphate, +S= high sulphate +Pi= plant colonized with *S. indica*, –Pi= non-colonized plant). Different asterisk indicates significant differences, * significant (p≤0.05), **(p≤0.01), ***highly significant (p≤0.001). Tukey test was used to check the significance. Data were analyzed three independent times in triplicates.

### Expression analysis of sulphate assimilation pathway genes of *S. indica* grown axenically and during colonization with the host plant

The up and down-regulation of sulfur assimilation genes were observed in case of WT and KD-*PiSulT*-*S. indica* that was grown axenically under low sulphate condition (**Supplemental Figure 13; Supplemental Table 4**). In case of WT *S. indica*, out of 22 selected genes from sulphate assimilation pathway, 11 genes; siroheme synthase (PiMET1), sulfite reductase cys-4 (PiMET5), sirohydrochlorin ferrochelatase (PiMET8), methylenetetrahydrofolate reductase (PiMET12), adenylylsulphate kinase (PiMET14), 3’-phospho-adenylylsulphate reductase (PiMET16), O-acetyl homoserine (thiol)-lyase (PiMET17), bisphosphate-3’-nucleotidase (PiMET22), sulphate adenylyltransferase (PiATPS), cystathione gamma-lyase (PiCYS3) and sulphate transporter (PiSulT) were found to be up-regulated. Amongst two genes i.e., PiSulT and PiMT16 were found to be 14 and 9.7 fold upregulated. In the case of KD-*PiSulT*-*P.indica*, 6 genes were found to be up-regulated i.e., transcription factor (PiMET4), 22-fold, methylinetetrahydrofolate reductase (PiMET12), 5.1-fold, methylenetetrahydrofolate reductase (PiMET13), 10-fold, adenylyl-sulphate kinase (PiMET14) 10.7-fold and sulphate transporter (*PiSulT*) 4.1-fold. A very high i.e., 48.6-fold up-regulation was found in the case of cyctathionine gamma-lyase (PiCYS3) **(Supplemental Figure 14; Supplemental Table 4)**. Only 3 genes i.e., siroheme synthase (PiMET1), 3’-phospho-adenylylsulphate reductase (PiMET16), and PiSulT were found to be up-regulated during colonization of the WT *S. indica* with the maize plant grown under low sulphate condition. A maximum of 12-fold up-regulation was found in the case of *PiSulT* under similar conditions (**Supplemental Figure 15; Supplemental Table 4)**. When KD-*PiSulT-S. indica* was colonized with the maize plants, 6 genes, i.e., sulfite reductase flavin-binding alpha-subunit (PiMET10), methylinetetrahydrofolate reductase (PiMET12), methylenetetrahydrofolate reductase (PiMET13) and sulphate adenylyltransferase (PiATPS), were found to be up-regulated and a maximum 24-fold, up-regulation was found in case of PiMET1. Interestingly, *PiSulT* was found to be down-regulated under similar condition as compared to WT ***S. indica* (Supplemental Figure 16; Supplemental Table 4**).

### Expression analysis of sulphate assimilation pathway genes of maize plants during colonization with WT and KD-*PiSulT*-*S. indica*

In case of maize plants colonized with WT *S. indica* and grown under low sulphate condition, out of 31 selected genes of sulphate assimilation pathway, only 3 genes i.e., Methylthioadenosine nuclease (ZmMTN), serineacetyltransferase2 (ZmSAT2) and sulfotransferase (ZmSOT) were found to be up-regulated by 2-5-fold (**Supplemental Figure 17; Supplemental Table 5)**. In the case of plants colonized with the KD-*PiSulT-S. indica*, 13 genes were found to be up-regulated. A maximum of 6.6 and 6.2-fold up-regulation was found in the case of serine acetyltransferase1 (ZmSAT1) and gamma-glutamyltransferase1 (ZmGGT1), respectively (**Supplemental Figure 18; Supplemental Table 5)**. We have also found the up-regulation of APS kinase (ZmAPSK). Interestingly, all three sulphate transporter genes; sulphate transporter1 (ZmST1), sulphate transporter (ZmST3.4) and sulphate transporter (ZmST4.1) were found to be down-regulated in plants either colonized with the WT or KD-*PiSulT-P.indica* (**Supplemental Figure 17 and 18; Supplemental Table 5).**

## DISCUSSION

The rhizosphere is observed as a hot spot for microbial activity. Microbes like bacteria, saprophytes, and mycorrhizal fungi, helps in nutrient enrichment for plants by mobilization and cycling of nutrients. Due to leaching, high reactive nature, complex forms and insolubility, many nutrients are unavailable to the plants. Hence, plants have developed many strategies to cope with nutrients deficiency including sulphur. For example, modulation of the root system architecture such as root length, modulation of transport activity with distinct transport affinities, substrate specificities to ensure appropriate flux (Aibara and Miwa, 2014). Low availability of nutrients like sulphate, phosphate and iron results in the less crop yields all over the world. Amongst nutrients sulphate is also play important role in the plant growth and development and its deficiency also causes less crop production. It has been reported that soil texture and rain are the major factors that affect sulfur availability in the soil. Sandy and silty soil have less organic matter and often low in sulfur because a high rainfall leaches out sulphate very easily from the root zone (Scherer, 2009). Sulphate is the main sulfur source for plants contributing to about 5% of total soil sulfur. Generally, more than 95% of soil sulfur are organically bounded (sulphate ester and carbon-bonded) and thus are not directly available for plants (Fitzgerald, 1976; Tabatabai, 1986; Leustek, 1996; Scherer, 2001; Scherer, 2009; Gahan and Schmalenberger, 2014). It has been observed that plants require sulfur concentration between 0.1 to 0.5% of dry weight and concentration below 0.1% is critical for the normal plant growth (Daigger and Fox, 1971; Kang and Osiname, 1976; Kamprath and Jones, 1986; Sakal et al., 2000; Marschner, 2011; Sutar et al., 2017). The optimum range of soil sulphate content considers from 0.3% to 1.0% of the dry weight of the soil (Little and Nair, 2009). A concentration of 3-5 ppm of sulphate in the soil is adequate for the growth of many plant species including maize (Sutar et al., 2017). Most of the sulphur in soil is present in an organically bounded form, which is released by the enzymatic action of bacteria and fungi, therefore, becomes available for utilization by plants (Kertesz and Mirleau, 2004; Gahan and Schmalenberger, 2014; Speck, 2015; Jacoby et al., 2017). Due to the low availability, sulfur is the limiting factor for crop production throughout the world and is a major challenge for agriculture. It has been reported that induction of the sulphate sensing and expression of high-affinity sulphate transporter is the key mechanism to increase the sulphate uptake rate in roots under low sulphate conditions (Takahashi, 2019). All the above strategy works only when plant roots are in contact with the nutrients in the rhizosphere. But in case of nutrient-depleted soil, another important strategy of plant-fungal association can work to take nutrients from nutrients deprive soil with the help of fungal nutrient transporter. In this study, functional characterization of a high-affinity sulphate transporter from *S. indica* and its role in the transfer of sulphate to the host plant has been demonstrated and how *PiSulT* is helpful to the plant growth and development under low sulphate condition has been established.

We found that PiSulT showed the highest similarity with the sulphate permease of *S. vermifera* and *Rhizoctonia solani* among the fungus group. Phylogenetic analysis indicates that PiSulT is more closely related to high-affinity sulphate transporter of fungi. Functional domains/motifs analysis of PiSulT polypeptide indicates that it is a sulphate transporter membrane protein having a typical C-terminal STAS domain, a signature motif of sulphate transporter “GLY” and a catalytic domain which is essential for a protein to be a sulphate transporter. It has been reported that in eukaryotes STAS domain plays an important role in sulphate transport and post-transcriptional regulation (Rouached et al., 2005; Yoshimoto et al., 2007). Additionally, sulphate permease signature motif was also identified in case of PiSulT polypeptide, the similar motif was also reported in case of sulphate transporter of fungus and plant thus support our data (Sandal and Marcker, 1994; Smith et al., 1995; Van De Kamp et al., 1999). We have observed similar percent colonization of *S. indica* in case of maize plant either grown under low or high sulphate condition which suggests that colonization is not dependent on the sulfur availability. Further, it was found that the colonization pattern of *S. indica* with a host plant was associated with the developmental stage of root tissue. We found that maturation and differentiation zone of lateral roots was densely colonized in comparison to the distal part of the apical root meristem. A similar pattern of colonization was also observed in the case of *S. indica* colonized with barley plants (Deshmukh et al., 2006), hence support our data.

A high transcript level of *PiSulT* under low sulfate concentrations (≤ 100 µm) indicates the high-affinity nature. Similar observations were also made in case of high-affinity phosphate transporter of algae, fungi and sulphate transporter of *S. cerevisiae*, therefore, these studies support our data (Chung et al., 2003; Yadav, 2010; Kankipati et al., 2015).

As our growth analysis showed that *PiSulT* complemented mutant and WT have similar and typical diauxic growth pattern which suggests that *PiSulT* efficiently complements the mutant. Previously, it has been shown that chromate enters the cells mainly through sulphate transporter and competitively inhibits sulphate uptake (Pereira et al., 2008). Several studies have been reported for uptake of chromate by sulphate transporter in different organism like in yeast, fungi, bacteria, and in mammalian cells (Ohta et al., 1971; Roberts and Marzluf, 1971; Campbell et al., 1981; Smith et al., 1995a; Cherest et al., 1997). Transport of sulphate and chromate is a type of competitive transport, and it depends on the concentration of either of the substrate. For this purpose, chromate toxicity was analyzed in the presence of different concentrations of sulphate. We observed that mutant (HK14) grew well and showed resistant to chromate as there was no uptake of chromate due to the absence of sulphate transporter gene, however, a very less growth was observed in case of WT and mutant complemented with the *PiSulT* (due to the uptake of chromate), therefore both were found to be susceptible to chromate toxicity. This observation confirms that *PiSulT* is a sulphate transporter.

Kinetics data reveals that *PiSulT* follows typical Michaelis-Menten kinetics. The apparent *Km* value was found to be 15 µM. Kinetic analysis of the sulphate uptake isotherm obtained in a range of low external sulphate concentrations (0–100µM) revealed that PiSulT has a high affinity for sulphate similar to those of other high-affinity sulphate transporters having *Km* values ranges from 4-14 µM (Smith et al., 1995b; Smith et al., 1995; Cherest et al., 1997; Smith et al., 1997; Takahashi et al., 2000; Vidmar et al., 2000; Yoshimoto et al., 2002; Howarth et al., 2003; Nocito et al., 2006). To best of our knowledge high-affinity sulphate transporter has *K*m ∼12 μM and low-affinity has a *K*m ∼100μM (Piłsyk and Paszewski, 2009). Previously, two high-affinity sulphate transporter of *S. cerevisiae*, have been reported Sul1 & Sul2 with a K*m* range between 4 to 10 µM (Smith et al., 1995a; Cherest et al., 1997). Therefore, we suggest that *PiSulT* belongs to high-affinity sulphate transporters. Our pH dependence analysis suggests that *PiSulT* functions maximum at pH 5. The effect of pH on sulphate uptake suggests that PiSulT transport might be facilitated by the proton gradient across the plasma membrane.

In order to know the role of *PiSulT* in sulphate transfer to the host plant, KD-*PiSulT*-*P.indica* were colonized with the host plant, this results in the reduction of the transfer of sulphate to the colonized plants as compared to the plants colonized with the WT *S. indica*, which suggests that *PiSulT* play a role in sulphate transfer from soil to host plant. A significantly higher expression of *PiSulT* in external hyphae than internal hyphae was observed, which indicates that external hyphae are the main site of *PiSulT* expression, and it is helping in the uptake of sulphate (available outside) to plant roots. A similar finding was also observed in case of *PiPT* when maize plants were colonized with the *S. indica*, this authenticates our data (Yadav, 2010). This expression pattern of *PiSulT* suggests that *S. indica* is supportive in the acquisition of sulphate from deprive range of sulphate concentration in soil rhizosphere with the help of hyphae.

Our study highlights the importance of *PiSulT* in the improvement of sulfur nutrition of the host plant particularly below plant’s required sulphate concentration (0.1 to 0.5% of plant dry weight) and below normal range of soil sulphate concentration (0.3% to 1.0% of dry weight of soil) for adequate plant growth and development. We observed that in the case of WT *S. indica* colonized plant biomass was 2.4 and 1.8-fold more than that of the non-colonized plants as well as from the KD-*PiSulT*-*P.indica* colonized plants, respectively. Importantly, it was found that the sulphate has an impact on the biomass of the maize plant colonized with *S. indica*. In our study, total sulfur content and biomass were found higher in the plants colonized with WT *S. indica* as compared with non-colonized and KD*-PiSulT-P.indica-*colonized plants, this suggests that sulphate is playing a role in augmenting plant yield or biomass, and this increase in the biomass is due to the *PiSulT*. Further, the effect of KD-*PiSulT-S. indica* colonization on plant metabolite content shows that low sulphate availability results in the low level of metabolites and sulfur-containing compounds such as glutathione. In a study, it has been shown that sulphate starvation affects other metabolic pathways in a pleiotropic manner (Nikiforova et al., 2005; Sieh et al., 2013). Our study demonstrates improved sulphur nutrition and the importance of *S. indica* in sulphur uptake for plant metabolism. It is interesting to note that the biomass promoting activity of *S. indica* was more under low sulphate condition as compared to that of sulphate-rich condition. A similar result was also found in the case of sulphate transporter of *M. truncatula* associated with mycorrhizal fungi (Sieh et al., 2013), hence validate our data.

Sulphate transport is the initial step in the acquisition and assimilation of sulfur and the flux of sulfur through the assimilatory pathway is likely to be linked to the regulation of sulphate transporters. Transcriptional regulation of several genes encoding enzymes of the sulfur assimilatory pathway in response to the plant sulfur status has also been reported (Droux, 2004; Casieri et al., 2012). Our data suggest that the expression of sulfur assimilation genes is dependent on the availability of sulphate. As we found more sulfur assimilation pathway genes were up-regulated during the KD-*PiSulT*-*S. indica* colonization with the plant, maybe these genes are helping the plant in getting more sulphate from the soil. The assimilation of sulphate occurs through a pathway that includes its uptake by specific permeases, activation of intermediates by ATP-dependent adenylation and reduction to sulfite and further to sulfide. Sulphate assimilation pathway genes particularly biosynthesis of methionine, cysteine, and S-adenosyl-methionine (SAM) were found to be up-regulated in WT axenically grown *S. indica* which indicates the active involvement of these genes in sulfur assimilation. The genes which are showing up-regulation in the case of KD-*PiSulT*-*P.indica* (grown axenically) indicate that they are responsive to external sulfur concentration and most probably involved in signaling under low sulphate condition. We conclude that starvation of any one of several amino acids results in the increased expression of genes encoding enzymes of multiple amino acid biosynthetic pathways. The down-regulation of genes in the case of KD-*PiSulT-S. indica* can be explained by the effects of reduced sulfur availability on the biosynthesis of amino acids, proteins, and sulfolipids.

In our study, many sulfur assimilation pathway genes appeared to be differentially expressed upon sulfur deficiency between non-colonized and colonized plants. In the case of maize plants colonized with WT *S. indica* and grown under low sulphate condition, out of 31 selected sulphate assimilation pathway-related genes, only 3 genes were found to be up-regulated as compared to 13 genes of plants colonized with the KD-*PiSulT-S. indica*. It is established that plants respond to a limited sulphate supply by increasing the expression of genes involved in the uptake and assimilation pathway. Amongst, 13 genes of a plant, we have found increased expression of APR, APS kinase and SAT. The increased expression of APR and ATP-sulfurylase indicates adapting response in the plant through the assimilatory pathway during the low supply of the sulphate by the fungus to the plant. We hypothesize that when KD-*PiSulT-P.indica* provides less sulphate to the plant during the colonization, in order to fulfill the high demand of sulphate in the fast-growing roots, programmed responses got induced. Increased expression of APR, APS kinase and SAT genes have been reported in case of maize plant under sulphate limited conditions and authors have suggested that these enzymes are important regulatory components of the sulfur assimilation pathway and got induced under sulphate-limited condition, therefore, helps the plant during the sulphate deficiency, thus these reports support our data (Hopkins et al., 2004). Roots growing in the soil may encounter glutathione originating from microbial activity or released by organic matter. This exogenous glutathione pool may be a valuable source of reduced sulphur for the fungi. It has been proposed that glutathione may play a role in the mycorrhizal symbiosis. Recently, it was suggested that mycorrhiza can improve sulphate availability to the root cell by extending their exploration horizon and, in exchange, receive reduced sulphur compounds from the root cell (Mansouri-Bauly et al., 2006). GGT is known to promote the hydrolysis of extracellular glutathione. As in our study, we found gamma-glutamyltransferase (ZmGGT1) highly up-regulated during the interaction of KD-*PiSulT-S. indica* and maize plant which suggests that glutathione will be available to the fungi in low quantity. We hypothesize that as the plant is not getting an adequate supply of the sulphate, therefore not giving the reduced sulphur to the fungi in exchange by increasing the expression of the ZmGGT1.Glutathione S-transferase (GST) has been proposed to play an integral role in the plant defense against the toxins and found to be induced during the chemical treatments and environmental stresses (Leustek et al., 2000). We assume that in this case also when there is less supply of the sulphate to the host plant by the KD-*PiSulT*-*S. indica*, GST got induced to generate a plant defense mechanism to avoid any harsh conditions. All other genes of sulfur assimilation pathway which were found up-regulated in case of KD-*PiSulT-P.indica* colonization with the plant, which is working either as a precursor/ regulatory component in the synthesis of amino acids or are involved as an intermediate in the sulfur assimilation may be helping the plant to get more sulphate during deficiency therefore it needs warrant investigation.

Our study also provides a new prospect to understand the sulphate transport network with and without a host plant. Additionally, the expression of the sulfur assimilatory pathway genes of the maize plant and their significance in balancing sulfur flux to sulphate demand of the plant for growth and development during interaction with the fungal partner will provide the insights into the sulphate management by the plant during sulphate deficiency. Thus, we suggest that in future *S. indica* can be used in the sulphate-deficient agriculture field to improve plant productivity.

## METHODS

### Plant, Fungi, Bacteria, and Yeast Strains

*Zea mays* (HQPM-5) plant and fungus *S. indica* were used throughout the study. *E. coli* XL1-Blue and DH5α were used for cloning purposes (Sambrook et al., 1989). Yeast sulphate transporter (Δsul1&Δsul2) double mutant HK14 (MATα *sul1*::*KanMX sul2::KanMX his3*Δ *leu2*Δ *lys2*Δ *ura3*Δ) and WT *S. cerevisiae* (BY4742) (MATα his3Δ leu2Δ lys2Δ ura3Δ) were used for complementation and kinetics (Kankipati et al., 2015) **(Supplemental Table 6)**. Both strains have the same BY background. Maize seeds were surface sterilized for 2 min in ethanol followed by 10 min in a NaClO solution (0.75% Cl) and finally washed six times with sterile water. Additionally, seeds were also treated with double-distilled H_2_O at 60°C for 5 min and were germinated on water agar plates (0.8% Bacto Agar, Difco, Detroit, MI) at 25 °C in the dark (Varma et al., 1999). Plants were grown under controlled conditions in a greenhouse with an 8 hours light (1000 Lux)/ 16 hours dark period at a temperature of 28°C with a relative humidity 60–70%. Surface sterilized pre-germinated maize seedlings were placed in pots filled with a mixture of sterile sand and soil in the ratio of 3:1 (garden soil from Jawaharlal Nehru University campus and acid-washed riverbed sand). Plants were weekly supplied with half-strength modified Hoagland solution containing the following: 5 mM KNO_3_, 5 mM Ca(NO_3_)_2_, 2 mM MgSO_4_, 10 μM KH_2_PO_4_, 10 μM MgCl_2_, 4 μM ZnSO_4_, 1 μM CaSO_4_, 1 μM NaMoO_4_, 50 μM H_3_BO_3_. Plant roots were harvested at different time points after inoculation and were assessed for colonization. To study colonization, ten root samples were selected randomly. Samples were softened in 10% KOH solution for 15 min and acidified with 1 N HCl for 10 min and finally stained with 0.02% Trypan blue (Phillips and Hayman, 1970; Dickson, 1998; Kumar et al., 2009). After 2 hours, samples were de-stained with 50% Lactophenol for 1–2 hours before observation under a light microscope (Leica Microscope, Type 020-518.500, Germany and Nikon Eclipse Ti). The distribution of chlamydospores within the root was taken as an index for studying colonization. Percent colonization was calculated for the inoculated plants according to the method described previously (McGonigle et al., 1990; Kumar et al., 2009).

### Isolation of *PiSulT* cDNA

Total RNA was isolated from *S. indica* grown axenically in KF medium (Hill, 2001) containing low sulphate (10µM) concentration with Trizol reagent (Invitrogen, USA) and cDNA was synthesized using a cDNA synthesis kit (Stratagene). *PiSulT* ORF (2292bp) was PCR amplified by using gene-specific primers (**Supplemental Table 7**). For directional cloning BamHI and XbaI sites were added in gene-specific forward and reverse primers respectively (**Supplemental Table 7**). For PCR reaction, *S. indica* cDNA was used as a template. PCR reactions were carried out in a final volume of 50 μl, containing 10 mM Tris-HCl (pH 8.3); 50 mM KCl; 1.5 mM MgCl2; 200 μM of dNTPs; 3 μM of each primer; 3 units of Phusion High-Fidelity DNA polymerase (Thermo Fisher Scientific and 60-100 ng of cDNA as template. PCR program was used as follows: 94°C for 2 min (1 cycle), 94°C for 45 sec, 60°C for 1 min 15 sec, 72°C for 1-2 min (35 cycles) and 72°C for 5 min (1 cycle). The PCR product was cloned into a pJET1.2 cloning vector (Promega) and further subcloned into the pYES2 yeast shuttle vector between BamH1 and Xba1 sites.

### Quantitative RT-PCR analyses

*S. indica* culture was grown in MN medium. Following contents were used / liter (MgCl_2_, 731 mg; Ca(NO_3_)_2_.4H_2_O, 288mg; NaNO_3_, 80 mg; KCI, 65 mg; Glucose, 10 g; NaFeEDTA, 8 mg; KI, 0.75 mg; MnCI_2_.4H_2_O, 6 mg; Zn Acetate, 2.65 mg; H_3_BO_3_, 1.5 mg; CuCl_2_,0.13 mg; Na_2_MoO_4_.2H_2_O, 0.0024 mg; Glycine, 3 mg; Thiamine Hydrochloride, 0.1 mg; Pyridoxine Hydrochloride, 0.1 mg; Nicotinic Acid, 0.5 mg; Myoinositol, 50 mg; Na_2_SO_4_ as per need, pH 5.5) (Bécard and Fortin, 1988). Further, *S. indica* culture were harvested at different time points and total RNA was isolated. The first strand of cDNA was synthesized with the Superscript cDNA synthesis kit (Clontech) from 3g of total RNA and used as a template for PCR with gene-specific primers (**Supplemental Table 7**). The reaction mixture was heated at 95 °C for 20 min and then subjected to 40 PCR cycles of 95 °C for 3s, 65 °C for the 30s, and 72 °C for 20 s. The heat dissociation curves confirmed that a single PCR product was amplified. The melting temperatures were 60-65 °C for the PCR products of the *PiSulT. S. indica* translational elongation factor gene (*PiTef*) was used as a control (Yadav, 2010). The level of target mRNA, relative to the mean of the reference housekeeping gene, was calculated by the relative ΔΔ*C*t method as described by the manufacturer.

### Phylogenetic and homology analysis

The functional sites in *PiSulT* and their pattern were determined using the PROSITE database. For identification purposes, blastX algorithm (www.ncbi.nlm.nih.gov) was used. Sequence alignments were done with ClustalΩ and BLOSUM62 with a gap penalty of 10 for insertion and 5 for extension (Henikoff and Henikoff, 1992; Thompson et al., 1994). Phylogenetic and molecular evolutionary analyses of *S. indica* putative *PiSulT* were constructed using *MEGA* X with the neighbor-joining analysis examined by bootstrap testing with 1000 repeats (Kumar et al., 2018).

### Complementation assay and growth analysis

For this purpose, *S. cerevisiae* WT BY4742 and yeast high-affinity sulphate transporter mutant strain HK14 were used (Kankipati et al., 2015). HK14 was transformed with a recombinant pYES2 vector having *PiSulT* by LiCl-PEG method (Bun-Ya et al., 1991; Riesmeier et al., 1992; Gietz et al., 1995; Akum et al., 2015; Jogawat et al., 2016). Yeast cells were grown at 30°C on SD media containing 0.1 mM of sulphate as a sole source of sulfur in the presence of 2% glucose (non-inducing condition) and 2% galactose (inducing condition) as sole carbon source separately. Cells were suspended in sterile distilled water and cell density was adjusted to A_600_= 0.1, followed by serial dilutions of 1/10. HK14 cells transformed with empty vector were used as a control. The drop test was performed to check the complementation. 30 µl suspensions were plated on solidified agar plates containing glucose and galactose separately. Plates were incubated for 2-3 days and the growth pattern of cells was analyzed. For the growth pattern of all three strains WT, mutant HK14 complemented with *PiSulT* and mutant complemented with empty vector pYES2 were also analyzed separately in SD media supplemented with different sulphate concentrations. For this purpose, cells were starved for sulfur source and then transferred to medium containing different concentrations of sulphate as the sole source of sulfur. The flasks were kept at 30°C and OD_600_ was measured to observe a comparison between the WT and the transformed mutant strains. Experiments were carried out in triplicates and were repeated thrice.

### Sulphate uptake and kinetics assay

Yeast cells were grown up to OD*A*_600_ of 1.5-2 in selective SD synthetic medium (YNB media with 2% glucose, lacking uracil) at 30^0^C for two days at 220 rpm in a metabolic shaker (Infors, Switzerland). Exponentially grown cells were washed with autoclave ddH_2_O and then transferred to sulfur starvation media, containing 2% of galactose (pH 5, with 50mM MES-KOH) at 30^0^C, 220 rpm for two days. Cells were harvested and resuspended at a cell density of 60 mg (wet weight)/ml. To start sulphate uptake, 50 µl of cells (preincubated for 10 min at 30^0^C) were aliquoted and different concentration of sulphate (1µM, 2µM, 4 µM, 6 µM, 10 µM, 25 µM, 75 µM, and 100 µM) was used. For this purpose, 0.5mM [^35^S] sodium sulphate (specific activity of 2000 cpm/nmol or 0.9 Ci/mol of sodium sulphate) was used. After 4 min, uptake was stopped by adding 5 ml of ice-cold sulfur starvation media. Cells were recovered on a glass microfiber filter and washed three times with 5 ml of ice-cold sulphate starvation media by centrifugation at 5000g for 5min at 4^0^C. For the blanks, ice-cold sulphate starvation media was added before the addition of [^35^S] sodium sulphate, and the cells were immediately filtered and washed. Further, filters were transferred into scintillation vials containing 5 ml of scintillation cocktail ‘O’ (CDH) and the radioactivity was measured with a scintillation counter (Liquid Scintillation Analyzer TRI-CARB 2100TR; Packard). Uptake assay was performed at room temperature (25^0^C). Sulphate accumulation (in pmol) was measured by standard mathematical calculations to convert scintillation count to pmole. The amount of sulphate transported by control (background) was used to normalize the data. Transport data at 10µM concentration was used for plotting the uptake graph. The rate sulphate uptake was expressed as nmol.min^-1^ x (mg dry weight)^-1^ or pmol/min/A_650._ GraphPad Prism 6 was used to plot nonlinear regression for sulphate uptake rate. Experiments were repeated three times and each time three replicates were taken.

### Development of RNAi Cassette and knockdown *S. indica*

A 452 bp unique fragment of *PiSulT* (**Supplemental Figure 9 ii**) was selected using the BLAST tool and analyzed for its uniqueness and RNA 2^0^ structures. This unique fragment was amplified using the gene-specific primers (**Supplemental Table 7**) and cloned into a pGEM-T cloning vector and subsequently subcloned into the pRNAi vector at the unique EcoRV site (Hilbert et al., 2012). This construct was named as pRNAi-PiSulT. Empty pRNAi and pRNAi-PiSulT was transformed into the *S. indica* as described previously (Yadav, 2010). In brief, chlamydospores were harvested from 14 days old *S. indica* culture and were germinated under glucose nutrition. For the cell wall disruption, β-Glucuronidase enzyme (Sigma: *Helix pomatia*) was used. Linearized pRNAi-PiSulT (1µg) was transformed into *S. indica* using electroporation at 12.5 kV/cm, 25-microfarad capacitance and 5-ms pulse length. Four transformed colonies (TC1, TC2, TC3, and TC4) were selected after primary and secondary selection using KF media containing 100µM and 200 µM concentration of hygromycin respectively (**Supplemental Fig. 10A)**. The transformation was confirmed by PCR using hygromycin gene-specific primers and siRNA analysis (**Supplemental Table 7; Supplemental Figure 10C & D**). All four transformants were tested for the expression of the silenced gene (*PiSulT*) by q-RT-PCR as described previously (Jogawat et al., 2016). Transformants obtained were named as “KD-*PiSulT*-*P.indica*”.

### Northern blot analysis

Northern blot was performed for siRNA detection. For this purpose, total RNA was isolated from KD*-PiSulT-P*. *indica* from the TC1 (in duplicate) and WT *S. indica* by using TRIzol reagent and probe was prepared by end labeling of the *PiSulT* end labeling primer (5′GTAATATCGACACGACCG) using [γ -^32^P] ATP and polynucleotide kinase as per the instructions described in manual (Molecular Labeling and Detection, Fermentas). Hybridization and autoradiography were performed as described (Yadav, 2010). RNA was dissolved in diethyl pyrocarbonate (DEPC) water, heated to 65 °C for 5 min, and then kept on ice. To this, polyethylene glycol (molecular weight of 8000, Sigma) was added to a final concentration of 5% and NaCl to a final concentration of 0.5 M. After incubation on ice for 30 min, this mixture was centrifuged at 10,000X*g* for 10 min. The supernatant obtained was mixed with the three volumes of ethanol. To precipitate the RNA, this mixture was kept at −20 °C for at 2 h. To obtain low molecular weight RNAs, the mixture was centrifuged for 10 min at 10,000X*g.* The pellet obtained was dissolved in DEPC treated water and heated at 65 °C for 5 min. Further, one-third volume of 4X loading solution (2xTBE (1xTBE is 0.09 M Tris-borate, pH 8.0, and 0.002 M EDTA), 40% sucrose, and 0.1% bromphenol blue) was added before loading on 15% urea-PAGE in 1xTBE. The RNA samples were electrophoresed at 2.5 V/cm and then blotted to a Hybond N^+^ membrane (Amersham Biosciences), and UV cross-linked. The membrane was prehybridized in 50% formamide, 7% SDS, 50 mM NaHPO_4_/NaH_2_PO_4_, pH 7.0, 0.3 M NaCl, 5X Denhardt’s solution (1X Denhardt’s solution is 0.02% Ficoll, 0.02% polyvinyl pyrrolidone, and 0.02% bovine serum albumin), and 100 mg/ml sheared, denatured salmon sperm DNA at 37 °C for at least 3 h. The probe was prepared by labeling the small fragment of *PiSulT* gene using [γ-^32^P] ATP and polynucleotide kinase as per the instructions manual (Molecular Labeling and Detection, Fermentas) and was added to the pre-hybridization solution. The hybridization was performed at 37 °C overnight, and the membrane was subsequently washed at 37 °C in 2X SSC (1X SSC is 0.15 M NaCl and 0.015 M sodium citrate) and 0.2% SDS for 15 min twice. Final washing was given only with 2X SSC at room temperature for 10 min and autoradiography was done. DNA oligonucleotides 16 and 22 nucleotides (nt) were used as molecular size markers for siRNA analysis.

### Bi-compartment assay

A 6-cm Petri dish (compartment 2) placed inside a 15-cm Petri dish (compartment 1) for setup bi-compartment experiment to make a physical barrier between both compartments. *S. indica* was grown in compartment 2. Surface-sterilized maize seeds were placed in compartment 1 to grow plants. The leafy shoots protruded through a groove cut in the lid of each dish and were fixed in one position by wrapping a sterile non-absorbent cotton wool around the portion of the subtending rhizome as it passed through the groove. In both the compartments co-cultivation MN media was used. Three sets were prepared for the experiment **(a)** maize plants colonized with WT *S. indica* **(b)** maize plants colonized with *S. indica*-KD*-PiSulT*, and **(c)** maize plants are grown alone without *S. indica*. In all the cases 10 µM sulphate concentration was used in compartment 1 as well as in compartment 2. For sets “*a*” and “*b*” to establish colonization between maize roots (compartment 1) and *S. indica* (compartment 2) a connective bridge was made by placing a 4 to 5 cm long agar strip so that *S. indica* can cross into the compartment 1. In the case of set “*c*”, a connecting bridge was also made to check any transfer of radioactive sulphate from compartment 2 to 1 due to diffusion, and this set was used as a control. As the colonization develops extraradical hyphae proliferate in the medium surrounding the roots in compartment 1 where they ramify and later sporulate. After colonization establishment, the MN media in compartment 2 of all three sets replaced with fresh MN media containing 100 µM Sulphate and 1 µM of ^35^S (specific activity, 200 mCi/mmol). Radioactivity determines in all three sets by autoradiography, and the amount of ^35^S incorporated measured by a liquid scintillation analyzer (Packard). The experiment was conducted three times independently.

### Spatial expression analysis of *PiSulT*

To determine the *PiSulT* expression in external hyphae and in internal hyphae of the *S. indica* colonized maize plant root, relative quantitative RT-PCR was performed as described (Yadav, 2010). In brief, external hyphae projecting out from the surface of the colonized root were collected by forceps. Approximately 2 mg of hyphae were collected per sample. In the case of internal hyphae sample collection, first, external hyphae were removed using forceps and or brushed off with a paint brush. Small pieces (5–10 mm) of colonized root were collected. Colonization was also confirmed in these collected root pieces as described previously (Narayan et al., 2017). RNA was isolated from these two samples, and cDNA was synthesized with the Superscript cDNA synthesis kit (Clontech) and used as a template for PCR with gene-specific primers for *PiSulT* and *PiTef* gene (control) (**Supplemental Table 7**). Quantitative RT-PCR was performed as described in the previous section.

### Plant metabolite measurements

To determine total sulfur contents, plants were harvested and dried in an oven at 150^0^C and crushed to make the fine powder. This powder was used for Energy Dispersive X-ray Fluorescence (ED-XRF) (PANalytical Epsilon 5) for measuring the total sulphate contents (per gram of dry weight). For sulphate ions measurement, 50 mg of frozen plant material was homogenized in 1 ml of deionized water containing 20 mg of polyvinylpolypyrrolidone. The sample was incubated with constant shaking at 4°C for 2h, at 95°C for 15 min and centrifuged at 14000 g for 20 min. 200 µl of supernatant was used to analyze by high-performance liquid chromatography (HPLC) (Agilent Technologies, Santa Clara, CA, USA, 1260 series) as described (Sieh et al 2013). Glutathione was extracted from the maize plant tissue by grinding 100 mg of frozen material in 1 mL of 0.1M HCl. The extract was centrifuged at 20,000 g for 10 min to remove cell debris. The supernatant was used to measure the total glutathione content after reduction with dithiothreitol and subjected to HPLC analysis using the monobromobimane derivatization (Sieh et al. 2013).

### Expression analysis of sulphate assimilation pathway genes of *S. indica* and maize plant

To investigate the relative expression of sulphate assimilation related genes of *S. indica* during axenic and colonization with the host plant, total RNA was extracted from the non-colonized and colonized *S. indica* and maize plants under sulphate-limited and sulphate-rich conditions. *S. indica* mycelia were grown in KF media for 7 days in high sulphate condition (10 mM) and filtered in minimal media containing low sulphate (LS = 10 μM) and high sulphate (HS = 10 mM) further grown for 7 days. Different sulphate concentrations were given to acclimatized *S. indica* by adding 10 μM (LS) and 10 mM (HS) in MN medium (0.4 mM NaCl, 2.0 mM KH_2_PO_4_, 0.3 mM (NH_4_)_2_HPO_4_, 0.6 mM CaCl_2_, 0.6 mM MgSO_4_, 3.6 mM FeCl_3_, 0.2 mM Thiamine hydrochloride, 0.1% (w/v) Trypticase peptone, 1 % (w/v) Glucose, 5 % (w/v) Malt extract, 2 mM KCl, 1 mM H_3_BO_3_, 0.22 mM MnSO_4_.H_2_O, 0.08 mM ZnSO_4_, 0.021 mM CuSO_4_, pH 5.8). After 1 week of different sulphate concentration supply, the fungus was immediately harvested and frozen in liquid nitrogen. In the case of colonization, maize plants were submerged in MN media supplemented with low sulphate (LS = 10 μM) and high sulphate (HS = 10 mM) for 1 week and the samples were frozen immediately, and the total RNA was isolated. Sulphate assimilation pathway-related genes of *S. indica* (**Supplemental Table 4**) and maize plant (**Supplemental Table 5**) were explored by BLASTp search. Two-step Real time-PCR protocol was used in different conditions. Real time-PCR reactions were performed on an ABI 7500 Fast sequence detection system (Applied Biosystems, Life Technologies, USA). The following cycles were used in the ABI 7500 Fast system (96 wells plates): pre-incubation at 95°C for 5 min, denaturation 94°C for 10 sec (4.8C/s), annealing at 60°C for 10 sec (2.5°C/s), extension at 72°C for 10 sec (4.8°C/s), 40 cycles of amplification and final extension at 72°C for 3 min. The Ct values were automatically calculated, the transcript levels were normalized against *PiTef* expression in the case of *S. indica* (Kumar et al., 2009) and against Actin in the case of maize and the fold change was calculated based on the non-treated control. The fold change values were calculated using the expression, where ΔΔC^T^ represents ΔC^T^ condition of interest gene-ΔC^T^ control gene. The fold expression was calculated according to the 2^-ΔΔC^T method mentioned elsewhere (Livak and Schmittgen, 2001).

### Statistical Methods

The statistical analyses were performed with Microsoft Excel 2010 and GraphPad Prism 8. The significance of the study was calculated using one-way ANOVA.

## Accession Numbers

Sequence data of *S. indica* sulphate transporter (*PiSulT*) can be found in the Gene bank Database (https://www.ncbi.nlm.nih.gov/genbank/) under the following accession number: MG816118.1

## Supplemental Data

Supplemental Table 1. List of domain hits.

Supplemental Table 2. Homology between highly similar fungus species with putative PiSulT

Supplemental Table 3. Homology of *PiSulT* with another organism.

Supplemental Table 4. Fold change of sulfur assimilation pathway genes of *S. indica*.

Supplemental Table 5. Fold change of sulfur assimilation pathway genes of the maize plant.

Supplemental Table 6. List of strains, and plasmids used in this study.

Supplemental Table 7. List of oligonucleotides used in this study.

Supplemental Figure 1. P*i*SulT ORF sequence and codon wise predicted amino acid sequences.

Supplemental Figure 2. P*i*SulT ORF sequence.

Supplemental Figure 3. Schematic representation of genomic orientation, ORF and protein translation of putative *PiSulT* gene.

Supplemental Figure 4. Multiple sequence alignment analysis.

Supplemental Figure 5. Multiple sequence alignment analysis.

Supplemental Figure 6. Colonization pattern of *S. indica* in maize roots.

Supplemental Figure 7. Chromate toxicity in the presence of different concentrations of sulphate.

Supplemental Figure 8. Effect of different concentrations of chromate on sulphate transport in WT complemented and mutant of sulphate transporter.

Supplemental Figure 9. pRNAi yeast shuttle vector and RNAi insert map.

Supplemental Figure 10. Knockdown of *PiSulT* gene of *S. indica*.

Supplemental Figure 11. Growth analysis of pRNAi-*PiSulT* transformed *S. indica*.

Supplemental Figure 12. Bi-compartment Petri dish culture system to study the transport of radiolabeled (^35^S) sodium sulphate to maize plants via *S. indica*.

Supplemental Figure 13. Relative expression analysis of putative sulphate assimilation pathway genes of WT-*S. indica* grew axenically under low sulphate.

Supplemental Figure 14. Relative expression analysis of putative sulphate assimilation pathway genes of KD-*S. indica* grew axenically under low and high sulphate conditions.

Supplemental Figure 15. Relative expression analysis of putative sulphate assimilation pathway genes of WT-*S. indica* during the colonized stage with maize plants under low and high sulphate conditions.

Supplemental Figure 16. Relative expression analysis of putative sulphate assimilation pathway genes of KD-*S. indica* during the colonized stage with maize plants under low and high sulphate conditions.

Supplemental Figure 17. Expression analysis of sulphate assimilation genes of maize plant during WT-*S. indica* colonized stage under low and high sulphate conditions.

Supplemental Figure 18. Expression analysis of sulphate assimilation genes of maize plant during KD-*PiSulT*-*S. indica* colonized stage under low and high sulphate conditions.

## AUTHOR CONTRIBUTIONS

AKJ has initiated the project. OPN and NV have performed the experiments. OPN, AKJ, AJ, and MD have designed the experiments. Chemicals were provided by AKJ and MD. The project was supervised by AKJ and MD. MS is written by OPN and AKJ.

The authors declare no conflict of interest

## ACKNOWLEDGMENTS

We are very thankful to Prof. Johan M. Thevelein, Laboratory of Molecular Cell Biology, Institute of Botany and Microbiology, KU Leuven, Kasteelpark Leuven-Heverlee, Flanders, Belgium, for providing yeast sulphate transporter mutant (HK14) and WT strain (BY4742) for study. OPN is grateful to the Indian Council of Medical Research (ICMR), Government of India for its financial support. NV is thankful to Jawaharlal Nehru University for providing a research fellowship. We are also very thankful to Prof. Alga Zuccaro, Institute for Genetics, Cologne Biocenter University of Cologne, Germany, for providing the pRNAi vector. AKJ and MD are thankful to Jawaharlal Nehru University for providing DST-PURSE-II, UPOE-II, and UGC-Resource NET-working grant.

